# A Hierarchical Clustering Algorithm Based on Silhouette Index for Cancer Subtype Discovery from Omics Data

**DOI:** 10.1101/309716

**Authors:** N. Nidheesh, K.A. Abdul Nazeer, P.M. Ameer

**Affiliations:** Department of Computer Science and Engineering, National Institute of Technology Calicut, Kerala, India.; Department of Electronics and Communication Engineering, National Institute of Technology Calicut, Kerala, India.

**Keywords:** hierarchical agglomerative clustering, Silhouette index, cancer subtype discovery, multi omics data, automatic cluster number identification, cluster number estimation, parameterless clustering algorithm, cluster analysis, gene expression data

## Abstract

Cancer subtype discovery from *omics* data requires techniques to estimate the number of natural clusters in the data. Automatically estimating the number of clusters has been a challenging problem in Machine Learning. Using clustering algorithms together with internal cluster validity indexes have been a popular method of estimating the number of clusters in biomolecular data. We propose a Hierarchical Agglomerative Clustering algorithm, named *SilHAC*, which can automatically estimate the number of natural clusters and can find the associated clustering solution. *SilHAC* is parameterless. We also present two hybrids of *SilHAC* with *Spectral Clustering* and *K-Means* respectively as components. *SilHAC* and the hybrids could find reasonable estimates for the number of clusters and the associated clustering solution when applied to a collection of cancer gene expression datasets. The proposed methods are better alternatives to the ‘*clustering algorithm - internal cluster validity index*’ pipelines for estimating the number of natural clusters.

---

Cluster analysis is a fundamental data mining technique that is used in a wide variety of areas for data analysis, especially in the Biomedical domain. There are many clustering algorithms proposed in the literature. *Hierarchical Clustering* is a classical clustering method and has been found to be used extensively in the Biomedical literature. Cluster analysis of *omics* data such as gene expression data has been a specific area where Hierarchical Clustering has been quite popular. The major reasons for such popularity, according to [1], are: 1) ease of use, as they require the setting of just one parameter - the number of clusters (or the information about where to cut the cluster *dendrogram*), and 2) the availability of implementations as part of many data analysis software.

Setting appropriate values for parameters is a hard task in the case of many clustering algorithms. Classical clustering algorithms such as *K-Means* require the number of clusters to be given as input by the user. Finding an appropriate value for the number of clusters is not an easy task in general. Exploratory data analyses such as discovering novel subtypes of cancers from molecular data of cancer tissues, require methods to estimate the exact number of natural clusters in the data. The prominent clustering algorithms available today lack this facility.

## A. Limitations in estimating the true number of clusters

The task of estimating the right number of clusters from an input dataset, in general, is hard. A reason for this limitation is that, in most of the cases, we are dealing with samples from a population and we need to estimate the number of true clusters in the population. The quality of estimation heavily depends on how well the sample at hand resembles the underlying population in the distribution of the data points. For instance, in the case of cancer gene expression data, we have the expression profile of a limited number of tissue samples and the distribution of the underlying population is not known. Furthermore, the aptness of the estimated number of clusters may be context dependent. Consider Figure 1. Visibly there are four groups of data points named G1, G2, G3, and G4. Consider the following three estimations of the number of clusters from the dataset - 1) *Four clusters*: Each group forming a cluster. 2) *Three clusters*: Groups G2 and G3 together forming a cluster and G1 and G4 forming their own individual clusters. 3) *Two clusters*: Groups G2, G3 and G4 together forming a cluster and G1 forming its own cluster. None of these groupings can be treated superior to the others without having any additional information about the data. The goodness of an estimated number of clusters and the associated clustering solution is dependent on the context and the dataset.

**Fig. 1.**
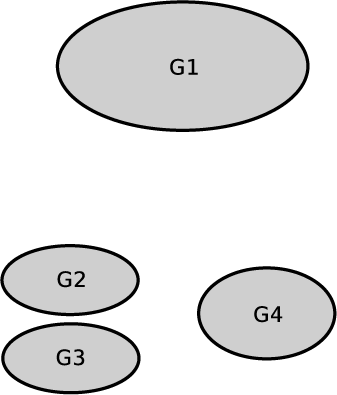
Issue in finding optimal number of clusters: Let G1, G2, G3 and G4 be four groups of data points. Three ways of inferring the number of clusters - 1) Four clusters: Each group forming a cluster. 2) Three clusters: Groups G2 and G3 together forming a cluster and G1 and G4 forming their own individual clusters. 3) Two clusters: Groups G2, G3 and G4 together forming a cluster and G1 forming its own cluster. The best clustering is context dependent.

The limitation discussed above is one among the inherent limitations that one faces when trying to estimate the optimal number of clusters from the data, and which are unavoidable. From within these limitations, the Biomedical research community has been using many methods to assess the optimal number of clusters from molecular data. A general technique followed is to use an *internal cluster validity index* in conjunction with a clustering algorithm (a ‘*clustering algorithm - internal cluster validity index*’ pipeline).

## B. Cluster validity index

The validity of clusters found out by a clustering algorithm is quantified using measures called *cluster validity indexes*. Cluster validity indexes are broadly classified into two: *Internal* and *External* [2]. External cluster validity indexes require the true class information of each data point along with the clustering result for the computation. In contrast, internal cluster validity indexes do not need true class information. Rather in general, they use just the clustering result and the similarity or dissimilarity measurements among the data points, to compute the index.

Several internal cluster validity indexes have been proposed in the literature. *Silhouette Index* [3] is one of the most successfully used cluster validity indexes in Biomedical data analysis. Details of computation of *Silhouette Index* are given in Section II. For a clustering solution of a dataset, the more the *average Silhouette Index* of all the data points closer to +1, the better the clustering solution is.

*Silhouette Index* has been widely used to make ‘*clustering algorithm - internal cluster validity index*’ pipelines. Such a pipeline is used to infer the optimal number of clusters in a dataset in the following way. Let *K-Means* be the clustering algorithm used to make the pipeline. *K-Means* is run a number of times with the value of the parameter *k* (which is the required number of clusters) varied from a minimum to a maximum (say, 2 to 20). For each of the resulting clustering solution, the *average Silhouette Index* is computed. The value of *k* that leads to a clustering solution which has the maximum *average Silhouette Index* is considered as an estimate of the number of natural clusters in the dataset.

## C. Cancer subtype discovery

Cancer is caused by faulty cellular mechanisms that lead to an out-of-control growth of tissues. There are several types of cancer. Each cancer type has multiple subtypes. A subtype may further have subtypes. Knowledge of the exact subtype of a cancer type helps in deciding the most appropriate treatment. *Cancer subtype discovery* is the task of finding out *previously unknown* subtypes [4].

Clustering algorithms have been instrumental in the discovery of cancer subtypes from *omics* data such as gene expression data.

## D. Silhouette Index in Biomedical literature

*Silhouette Index* has been well-received among Biomedical researchers. A recent study on the performance of Biomedical clustering algorithms [5], considers *Silhouette Index* as one of the best internal cluster validity index and suggests the use of *Silhouette Index* in the cluster analysis of Biomedical data where no ground truth class information of data points are available. *Silhouette Index* is found to be used for two purposes in the literature related to biomolecular data processing. One is as a measure to check the validity of obtained clustering solution. The other is to use along with a clustering algorithm to estimate the number of clusters in a dataset (such as in discovering cancer subtypes from *omics* data). Finding novel subtypes of cancers from *multi-omics* data is a key area of current Biomedical research. *Silhouette Index* has been in widespread use in estimating the number of natural clusters in the biomolecular data.

The paper [6] introduced a multi-view clustering approach (*CMC*) and its enhanced version (*ECMC*) for subtype identification from heterogeneous cancer datasets. The authors employed both *Spectral Clustering* and *K-Means* together with *Silhouette Index* to identify novel cancer subtypes. In [7], the authors presented an integrative clustering method for *multiomics* data of cancer tissues, named ‘*LRA Cluster*’, which employs ‘*K-Means-Silhouette Index pipeline*’ to determine the number of clusters (subtypes). The paper [8] presented a pattern fusion approach (PFA) for identifying cancer subtypes from heterogeneous *omics* data. The authors used *K-Means* to cluster the resulting data of the PFA process and considered the number of clusters indicated by *average Silhouette Index* as the number of subtypes. In [9], the authors proposed ‘*intNMF*’ for integrative clustering of *multi-omics* data of cancer tissues, to identify novel subtypes. The authors employed *average Silhouette Index* as one of the methods to get an optimum number of clusters. In [10], the authors proposed ‘*Integrated Consensus Clustering*’ for cancer subtype discovery by integrated analysis of *mRNA*, *miRNA* and *lncRNA* expression data. *Silhouette Index* is one of the methods they employed to find out the optimal number of clusters (subtypes). In [11] the authors proposed ‘*Similarity Network Fusion*’ (SNF) to fuse graph representation of cancer *multi-omics* data (obtained from TCGA) and applied *Spectral Clustering* together with *Silhouette Index* to get coherent clusters which represent subtypes.

## E. Our contributions

We propose a Hierarchical Agglomerative Clustering algorithm named *SilHAC* which uses a *Silhouette Index* based criterion for selecting the pair of clusters to merge, in the iterative merging process for making the cluster hierarchy. A salient feature of *SilHAC* is that it can automatically find an optimal number of clusters and find the associated clustering solution without taking any input other than the dataset.

We also propose two hybrid versions of *SilHAC*. A major goal of the hybrids is to minimize execution time. The hybrids employ a clustering algorithm to reduce the number of initial clusters to deal with by the iterative cluster merging process of *SilHAC*. We selected *Spectral Clustering* and *K-Means* to build the hybrids as these two algorithms along with *Silhouette Index* have been widely used in Biomedical literature as a method to get optimal number of clusters and associated clustering solution. However, one can use any other clustering algorithm to build a hybrid.

## I. RELATED WORK

Hierarchical Agglomerative Clustering (HAC) is a classical clustering algorithm. The algorithm works as follows. Initially the algorithm considers each data point belonging to its own cluster. At each step, the algorithm merges the most appropriate pair of clusters. The iterative merging process continues until there is only one cluster. The appropriateness of a pair for merging is decided by a criterion. HAC has many variants which differ from one another in the merge criterion employed.

Ward proposed a framework for HAC in [12]. According to the framework, one can employ an *objective function* that reflects the chosen criterion for selecting the pair of clusters for merging. The pair which leads to the maximal value for the objective function is selected for merging at each step. Ward suggested using the *total within-cluster sum of squared errors (TWSSE*) as an objective function.

Initially, as each data point is considered belonging to its own cluster, the cluster mean point coincides with the data point of the cluster for each cluster. So the *TWSSE* at this stage is *zero*. *TWSSE* increases at each subsequent steps when two clusters are merged (unless the data points of the merged clusters coincide). At each step, the algorithm selects the pair of clusters, which when merged, leads to the minimum increase in the *TWSSE*, for actual merging.

The variant of HAC with *TWSSE* as objective function (to minimize) is popularly known in the literature both as *Hierarchical Agglomerative Clustering (Ward’s method*) and as *Hierarchical Agglomerative Clustering (Ward’s minimum variance method*).

*SilHAC* differs from *Hierarchical Agglomerative Clustering* (*Ward’s method*) in many ways. *SilHAC* straightforwardly gives an optimal number of natural clusters in the dataset (and the associated clustering solution). *SilHAC* employs *average Silhouette Index* as the objective function (to maximize) for selecting clusters to merge whereas *Ward’s method* uses the *TWSEE* as the objective function (to minimize). The proposed hybrids of *SilHAC* start the iterative merging of clusters from the clusters produced by algorithms such as *K-Means* and *Spectral Clustering* whereas *Ward’s method* and other variants of HAC starts iterative merging from singleton clusters formed by each data point.

Another method related to the proposed methods is *Cross-Clustering* (*CC*) presented in [13]. *CC* is a combination of two well known Hierarchical Agglomerative Clustering algorithms - *Ward’s minimum variance* and the *Complete Linkage*. Similar to *SilHAC*, *CC* can automatically estimate the number of natural clusters in a dataset and can find the associated clustering solution.

A key difference of *CC* from *SilHAC* and its hybrids is that *CC* is a *partial clustering algorithm* (*CC* does not give the cluster membership information for all the data points of the input dataset) whereas *SilHAC* and its hybrids are *full clustering algorithms*.

## II. MATERIALS AND METHODS

### A. The proposed method

The basic idea of *SilHAC* is to use *Average Silhouette Index* as the objective function in *Ward’s framework*. For the hybrids, unlike Ward’s algorithm, the agglomerative merging of clusters starts from a set of clusters formed using the component clustering algorithm. We present a single algorithm for both *SilHAC* and the hybrids. The essence of the proposed method is given as Algorithm 1. The algorithm works in *SilHAC mode* or the *hybrid mode* depending on the value of a parameter named ‘*limit*’. If *limit* has a value greater than or equal to the size of the input dataset, the algorithm runs in *SilHAC mode*. Otherwise, it runs in the *hybrid mode* where the component clustering algorithm (*K-Means* or *Spectral Clustering*) is invoked first to get ‘*limit*’ number of clusters for *SilHAC* to begin its hierarchical merging process.

The relevant definitions of terms and functions used in Algorithm 1 are explained below.

#### 1) Definitions

##### Clustering Vector

*SilHAC* and the hybrids return the clustering result as a *clustering vector*. A *clustering vector* is a vector which indicates the cluster to which each data point belongs. Its size is equal to the number of data points in the input dataset. For instance, if we have a set of *five* data points with the first *three* data points belonging to cluster number 1 and the rest belonging to cluster number 2, then the corresponding *clustering vector* will have component values in the following order: 1, 1, 1, 2, 2.

##### ComputeDissimilarityMatrix(D)

This function takes a dataset *D* as input and returns a dissimilarity matrix. If *M* is a dissimilarity matrix then *M* [*i, j*] = *M* [*j, i*] = the measure of dissimilarity between the data points *i* and *j*. Any suitable measure can be used as the measure of dissimilarity in practice. A commonly used measure of dissimilarity is the *Euclidean Distance* and hence, for our implementation, we considered the *Euclidean Distance*. Hereafter we use the terms *dissimilarity* and *distance* interchangeably.

##### GetInitialCV(D, limit)

This function (given as Algorithm 2) takes a dataset *D* and an integer value named ‘*limit*’ as inputs and returns an initial clustering vector for *SilHAC*’s iterative cluster merging process. The parameter *limit* indicates the upper limit on the number of initial clusters that the merging process of a hybrid of *SilHAC* deals with. When the number of data points is less than *limit*, the algorithm runs in pure *SilHAC* mode and otherwise in the hybrid mode. In the hybrid mode, the component clustering algorithm is applied to the dataset to get *limit* number of clusters. The sole purpose of the component clustering algorithm in the *SilHAC* hybrids is to reduce the number of initial clusters to deal with by *SilHAC* and in turn, to reduce the time of execution of *SilHAC* while dealing with large datasets.

###### Algorithm 1 *SilHAC* (The proposed method) in hybrid form. If *limit* < |D| the algorithm runs in *SilHAC mode*.

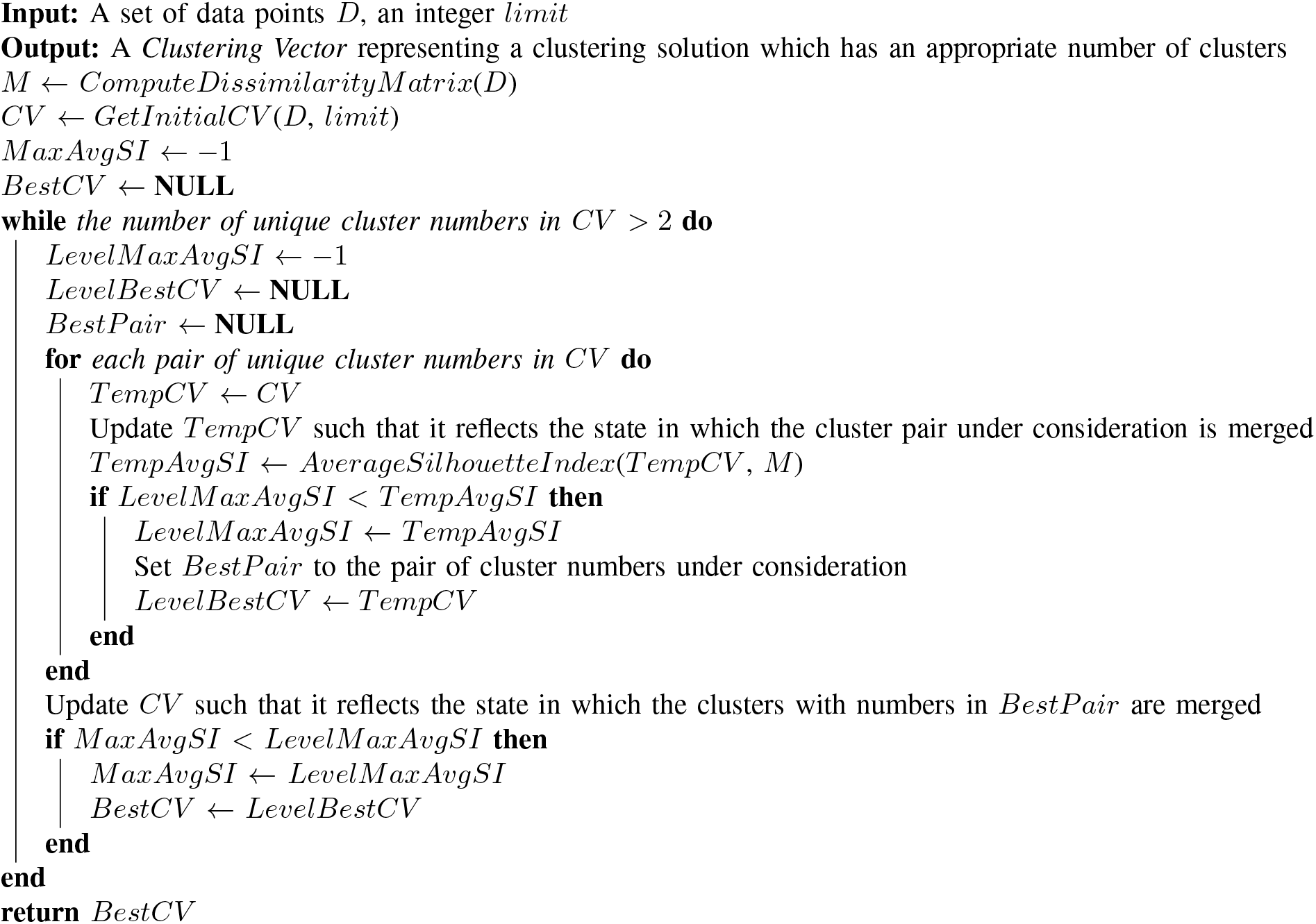

##### AverageSilhouetteIndex(CV, M)

This function computes the average of the *Silhouette Indexes* of all the data points according to the clustering solution given by the clustering vector *CV* and the distance matrix *M*.

*Silhouette Index* of a data point *i* is computed as follows [3].

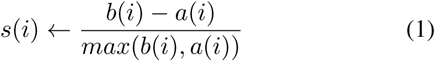

Here *a*(*i*) is the average of the distances of the data point *i* from every other data point of the cluster to which *i* belongs. Computation of *b*(*i*) is illustrated through the following example scenario. Let there be three clusters (say, *A*, *B* and *C*) in addition to the cluster of *i*. Let *bA*(*i*) be the average of the distances from *i* to all the data points in *A*. Similarly, let *bB*(*i*) and *bC*(*i*) be the average of the distances from *i* to all the data points in *B* and *C* respectively. Now, *b*(*i*) is the minimum among *bA*(*i*)*, bB*(*i*) and *bC*(*i*).

The *Silhouette Index* of a data point ranges from −1 to +1. Once we have the *Silhouette Indexes* of all the data points of the dataset computed, we can find the *average Silhouette Index* for the whole dataset. This value is returned by the function *AverageSilhouetteIndex*.

#### 2) The algorithm

The algorithm (Algorithm 1) works as follows. It takes a dataset *D*, and an integer *limit* as inputs. *limit* decides whether the algorithm runs in *SilHAC mode* or the *hybrid mode*. At first, a dissimilarity matrix for the data points is prepared which is necessary for computing *average Silhouette Index*. Then the algorithm prepares an *initial clustering vector*, depending on the value of the *limit* parameter, by calling the function *GetInitialCV*. This step is the only step where the component clustering algorithm is involved in a hybrid of *SilHAC*.

In essence, the rest of the algorithm does the following. It takes every pair of clusters and computes the *average Silhouette Index* for the clustering resulted by (temporarily) merging the pair. The pair which leads to the maximum *average Silhouette Index* (*BestPair*) is then (actually) merged to reduce the number of clusters by one. The *average Silhouette Index* of the resulting clustering solution and the corresponding clustering vector, are recorded after each (actual) merging. This procedure is followed until there are only two clusters left. The clustering vector which corresponds to the clustering solution which has the maximum *average Silhouette Index* is finally presented as the output by the algorithm.

##### Algorithm 2 *GetInitialCV*: This function prepares an initial clustering vector for *SilHAC*’s agglomerative cluster merging process.

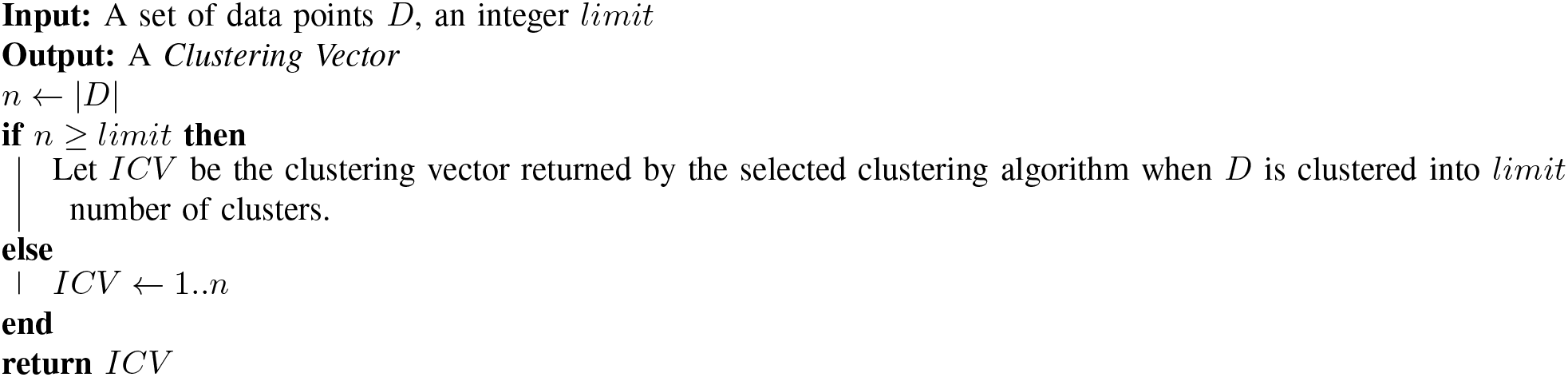

#### 3) Implementation

We selected *R* [14] for the implementation of *SilHAC* and the hybrids. The implementation has been made to look similar to the *R*’s native implementations of Hierarchical Agglomerative Clustering algorithms (provided by the *hclust* function of *R*). Our implementation facilitates *dendrogram* visualization. Moreover, the *R* function *cutree* can be applied on the resulting *dendrogram* to get any arbitrary number of clusters as we do with the results of the *hclust* function. For computing *Silhouette Index* we employed the *silhouette* function provided by the *cluster* package [15].

*Spectral Clustering* and *K-Means* are the two algorithms selected to build hybrids of *SilHAC*. For *Spectral Clustering* we employed the method *specc* of the *kernlab* package [16]. For *K-Means* we employed *R*’s native implementation of *Lloyd’s algorithm*.

### B. Datasets

Ten cancer gene expression datasets were used for analyzing the capability of the proposed methods. These datasets are the ones used in [17]. The selected group consists of *five* datasets from NCBI Gene Expression Omnibus (GEO) [18] and *five* datasets introduced in [19]. The datasets introduced in [19] have been used as a benchmark in many studies on clustering based identification of cancer subtypes.

The properties of the datasets are given in brief in Table I and Table II. The datasets with names GEO1, GEO2, GEO3, GEO4 and GEO5 of Table I are feature reduced (gene/probe reduced, as described in [17]) versions of the NCBI GEO datasets with accession numbers GSE51082 (GPL-97), GSE57162 [20], GSE66354 [21], GSE85217 [22] and GSE94601 [23] respectively. The names LEU, NOVA, STJ, LUNG and CNS represent the datasets *Lukemia*, *Novartis Multi-tissue*, *St. Jude Leukemia*, *Lung Cancer* and *Central Nervous System Tumors* respectively, of [19].

**TABLE I.** Cancer gene expression datasets from NCBI GEO: The datasets GEO1, GEO2, GEO3, GEO4 and GEO5 are feature reduced (gene/probe reduced, as described in [17]) versions of the NCBI GEO datasets with accession numbers GSE51082 (GPL-97), GSE57162, GSE66354, GSE85217 and GSE94601 respectively.

| Dataset name | Cancer type | No. of genes/probes | No. of samples | No. of samples (Subtype-wise) |
| --- | --- | --- | --- | --- |
| GEO1 | Leukemia | 2000 | 140 | 1) AML (37)<br>2) Chronic Lymphocytic Leukemia (41)<br>3) Chronic Myeloid Leukemia (22)<br>4) Myelodysplastic Syndrome (10)<br>5) Precursor B-cell ALL (17)<br>6) T-cell ALL (13) |
| GEO2 | Renal Cell Carcinoma (RCC) | 2000 | 191 | 1) Renal Oncocytoma (36)<br>2) Papillary RCC (57)<br>3) Chromophobe RCC (35)<br>4) Clear Cell RCC (63) |
| GEO3 | Brain Tumors | 2000 | 136 | 1) High Grade Glioma (21)<br>2) AT/RT (17)<br>3) Medulloblastoma (19)<br>4) Pilocytic Astrocytoma (15)<br>5) Posterior Fossa Ependymoma (55)<br>6) Supratentorial Ependymoma (9) |
| GEO4 | Medulloblastoma | 2000 | 763 | 1) Group 3 (144)<br>2) Group 4 (326)<br>3) SHH (223)<br>4) WNT (70) |
| GEO5 | Lung Carcinoma | 2000 | 137 | 1) Adenocarcinoma (73)<br>2) Large Cell Lung Cancer (29)<br>3) Large Cell-Neuroendocrine Carcinoma (11)<br>4) Small Cell Lung Cancer (3)<br>5) Squamous Cell Carcinoma (21) |

**TABLE II.** Cancer gene expression datasets taken from [19]:

| Dataset name | Cancer type | No. of genes/probes | No. of samples | No. of samples (Subtype-wise) |
| --- | --- | --- | --- | --- |
| LEU | Leukemia | 999 | 38 | 1) ALL-B(19)<br>2) ALL-T (8)<br>3) AML (11) |
| NOVA | Multiple cancers | 1000 | 103 | 1) Breast Cancer (26)<br>2) Prostate Cancer (26)<br>3) Lung Cancer (28)<br>4) Colorectal Cancer (23) |
| STJ | Pediatric Acute Leukemia | 985 | 248 | 1) BCR-ABL (15)<br>2) E2A-PBX1 (27)<br>3) "Hyperdiploid > 50" (64)<br>4) "MLL Rearrangement" (20)<br>5) ALL-T (43)<br>6) TEL-AML1 (79) |
| LUNG | Lung cancer | 1000 | 197 | 1) Adenocarcinoma (139)<br>2) Normal Lung Tissue (17)<br>3) Squamous Cell Carcinoma (21)<br>4) Carcinoid (20) |
| CNS | Central Nervous System tumors | 1000 | 42 | 1) Medulloblastoma (10)<br>2) AT/RT (10)<br>3) Malignant Glioma (10)<br>4) Normal Cerebellum Tissue (4)<br>5) PNET (8) |

### C. Setup for the comparative study

We conducted a comparative study of the performances of *SilHAC*, two hybrids of *SilHAC* and two traditional ‘*clustering algorithm - internal cluster validity index*’ pipelines, in identifying the optimal number of clusters in a dataset. The two hybrids of *SilHAC* considered are *SilHAC-Spectral Clustering hybrid* (hereafter referred to as *SilHAC-SC*) and *SilHAC-K-Means hybrid* (hereafter referred to as *SilHAC-KM*). Two traditional ‘*clustering algorithm - internal cluster validity index*’ pipelines used for the study are *Spectral Clustering - Silhouette Index pipeline* (hereafter referred to as *SC-SI*) and *K-means-Silhouette Index pipeline* (hereafter referred to as *KM-SI*).

For *SilHAC-SC*, *Spectral Clustering* was set to run 20 times (due to *Spectral Clustering*’s non-deterministic nature) to cluster the input dataset into 25 clusters. The best clustering solution (the solution which has the maximum *average Silhouette Index*) was then selected as the initial set of clusters for the iterative merging process.

Similarly for *SilHAC-KM*, *K-Means* was set to run 100 times (being non-deterministic) setting *k* = 25 to get 25 clusters. The best clustering solution (the one which has the minimum *total within-cluster sum of squared errors*) was then selected as the initial set of clusters for the agglomerative merging process.

For the *SC-SI* pipeline, *Spectral clustering* was set to run 20 times for each value of *k* in the range 2-20. For each *k*, the clustering solution which has the maximum *average Silhouette Index*) was selected. From among these 19 clustering solutions, the one with the maximum *average Silhouette Index* was then taken as the final clustering solution.

Similarly, for the *KM-SI* pipeline, *K-Means* was run 100 times for each value of *k* in the range 2-20. For each *k*, the clustering solution which has the minimum *total within-cluster sum of squared errors* was selected. From among these 19 clustering solutions, the one with the maximum *average Silhouette Index* was then taken as the final clustering solution.

All the datasets were subjected to *min-max normalization* before used for the analysis. The seed for the random number generator was set to *one* for the repeatability of the analyses.

## III. RESULTS AND DISCUSSION

The proposed methods performed well in finding reasonable estimates for the natural number of clusters and also in appropriately clustering the data points of the input datasets. A comparison of the proposed methods with that of the *two* selected ‘*clustering algorithm - Silhouette Index*’ pipelines is given in Table III. The values shown in the column *ASI* are the highest *Average Silhouette Indexes* (*ASI*) rounded off to *two* decimal points. In the case of each method and each dataset, if the difference of the *ASI* of a clustering solution from the highest ASI (when non-rounded) is less than or equal to 0.005, the *ASI* is treated as equal to the highest *ASI*. This is done to avoid the error in cluster number estimation which may occur due to rounding. It can be seen from the table that *SilHAC* and the hybrids provide promising results comparable to the popular *SC-SI* and *KM-SI* pipelines in finding reasonable estimates for the true number of clusters. Moreover, *SilHAC* and its hybrids had lower number of misclassifications for many of the datasets.

**TABLE III.** Performance comparison of the proposed methods (*SilHAC*, *SilHAC-SC* and *SilHAC-KM*) with that of two traditional *clustering algorithm - Silhouette Index* pipelines (*SC-SI* and *KM-SI*). *SilHAC-SC* stands for the hybrid of *SilHAC* and *Spectral Clustering*, with *Spectral Clustering* run 20 times for *k* = 25. *SilHAC-KM* stands for the hybrid of *SilHAC* and the standard *K-Means*, with *K-Means* run 100 times for *k* = 25. *SC-SI* and *KM-SI* stand for ‘*Spectral Clustering - Silhouette index*’ pipeline and ‘*K-Means - Silhouette index*’ pipeline respectively, with *Spectral Clustering* executed 20 times for each value of *k* in the range 2-20 and with *K-Means* executed 100 times for each value of *k* in the range 2-20. *ASI* stands for *average Silhouette Index*. When multiple clustering solutions resulted in the same highest *ASI*, the corresponding numbers of clusters are shown comma-separated. In each case, the best *ASI* and the number of clusters corresponding to the clustering with the best *ASI* are shown.

| Dataset | Ground truth |  | *SilHAC* |  | *SilHAC-SC* |  | *SilHAC-KM* |  | *SC-SI* |  | *KM-SI* |  |
| --- | --- | --- | --- | --- | --- | --- | --- | --- | --- | --- | --- | --- |
|  | Classes | ASI | Clusters | ASI | Clusters | ASI | Classes | ASI | Clusters | ASI | Clusters | ASI |
| GEO1 | 6 | 0.47 | 4 | 0.51 | 4 | 0.51 | 4 | 0.51 | 4 | 0.51 | 4 | 0.51 |
| GEO2 | 4 | 0.22 | 2 | 0.41 | 2 | 0.40 | 2 | 0.43 | 2 | 0.43 | 2 | 0.43 |
| GEO3 | 6 | 0.28 | 4, 5 | 0.32 | 4, 5 | 0.30 | 4 | 0.32 | 4 | 0.32 | 4 | 0.32 |
| GEO4 | 4 | 0.29 | 3 | 0.38 | 3 | 0.37 | 3 | 0.38 | 3 | 0.38 | 3 | 0.38 |
| GEO5 | 5 | 0.21 | 3, 4 | 0.25 | 3, 4 | 0.24 | 3, 4 | 0.23 | 3 | 0.24 | 3 | 0.25 |
| LEU | 3 | 0.17 | 3, 4, 5 | 0.17 | 3, 4, 5 | 0.17 | 3, 4 | 0.17 | 3 | 0.17 | 3 | 0.17 |
| NOVA | 4 | 0.26 | 2, 4 | 0.27 | 2, 4, 5 | 0.27 | 2, 4, 5 | 0.27 | 2, 4 | 0.27 | 2, 4 | 0.27 |
| STJ | 6 | 0.21 | 5, 6 | 0.21 | 5, 6 | 0.21 | 5, 6 | 0.21 | 5 | 0.21 | 6 | 0.22 |
| LUNG | 4 | 0.22 | 2 | 0.35 | 2 | 0.35 | 2 | 0.35 | 2 | 0.35 | 2 | 0.35 |
| CNS | 5 | 0.13 | 2 | 0.31 | 2 | 0.35 | 2 | 0.35 | 2 | 0.28 | 2, 3 | 0.28 |

In the case GEO1 dataset, all the methods are found to treat the true classes (subtypes) corresponding to *AML*, *Chronic Myeloid Leukemia* and *Precursor B-Cell ALL* as belonging to one cluster. The other *three* classes have been correctly identified. There were no other misclassifications for any of the methods.

For the GEO2 dataset, all the methods grouped the samples into *two* clusters in more or less the same fashion - one cluster comprising majority of *Chromophobe RCC* samples and the other comprising majority of the samples of the other *three* classes. *SilHAC* and *SilHAC-SC* had the minimum number of samples misclassified (*three* each).

*SilHAC* and *SilHAC-SC* methods could find *five* clusters in the case of GEO3 dataset whereas the other methods found only *four* clusters. GEO3 dataset has actually *six* classes (subtypes). While finding *five* clusters, *SilHAC* treated the classes of *High Grade Glioma* and *Pilocytic Astrocytoma* as one cluster and the other three classes were almost correctly identified (but suffering a misclassification of just one *Medulloblastoma* sample). *SilHAC-SC* also behaved in a similar fashion but suffered more misclassifications. *SilHAC-KM* also merged the above *two* classes. In addition, the method merged the classes corresponding to *Posterior Fossa Ependymoma* and *Supratentorial Ependymoma*. The traditional methods (*SC-SI* and *KM-SI* pipelines) had comparatively more number of misclassifications.

GEO4 is the biggest dataset considered for the study. It has 763 samples. For the dataset, all the methods found out *three* clusters. All the methods clubbed together the samples of *Group 3* and *Group 4* subtypes. Other than that, there were only a few (less than 10) misclassifications for each of the methods. *SilHAC-KM* had only *two* misclassifications.

For the GEO5 dataset which has *five* classes, *SilHAC* and its hybrids could identify *four* clusters whereas the others identified only *three* clusters. *SilHAC* and the hybrids merged the classes corresponding to *Large Cell Carcinoma* (*Large Cell Lung Cancer* and *Large Cell Neuroendocrine Carcinoma*) into one cluster.

For the *LEU* dataset (which has *three* classes), all the methods correctly identified the *three* classes with no misclassifications. *SilHAC* and the hybrids, in addition, identified *four* and *five* clusters for the datasets. When identified *four* clusters, *SilHAC* and its hybrids subdivided the *ALL-B* class (having 19 samples) into *two* clusters - one with 17 samples and the other with *two* samples. *SilHAC* and *SilHAC-SC* further divided the two-sample cluster mentioned above into *two* singleton clusters, when they identified *five* clusters.

For the *NOVA* dataset which has four classes, all the methods identified *two* and *four* clusters. All the methods except *SilHAC-KM* misclassified *two* samples each when they partitioned the dataset into *four* clusters. *SilHAC-KM* misclassified only *one* sample. *SilHAC-SC* and *SilHAC-KM* could also identify *five* clusters in the dataset in which case the additional cluster was constituted by *one* sample of *Breast Cancer*.

For the *STJ* dataset, which has *six* classes, all the methods except the *SC-SI* pipeline could estimate *six* as the optimal number of clusters. The number of samples misclassified ranged from *eight* to 14 with *KM-SI pipeline* having *eight* misclassifications. The *SC-SI* pipeline could identify only *five* clusters.

For the *LUNG* dataset, all the methods identified *two* clusters and all except *SilHAC-KM* produced the same clustering result - the class corresponding to *Carcinoid* subtype as the first cluster and the remaining *three* classes together forming the second cluster. *SilHAC-KM* had a few misclassifications.

For the *CNS* dataset, which has *five* classes, all the methods identified *two* clusters. The *KM-SI pipeline* in addition identified *three* clusters where it could correctly identify the class of *normal cerebellum tissues*. *SilHAC* perfectly delineated the class of *normal cerebellum tissues* from the classes of cancerous samples. Both *SilHAC-SC* and *SilHAC-KM* methods achieved the highest *average Silhouette Index* which is notably higher than those of *SC-SI* and *KM-SI* methods. The hybrids too grouped normal tissue samples as one cluster and all the cancerous samples as the other cluster, but by suffering the misclassification of one normal sample. The clusters obtained by *SC-SI* and *KM-SI* pipelines suffer more number of misclassifications.

An advantage of *SilHAC* and its hybrids, is that more reliable results are produced by them when compared to those of the traditional ‘*clustering algorithm - internal cluster validity index*’ pipelines constituted using non-deterministic clustering algorithms such as *K-Means* and *Spectral Clustering*. While clustering a dataset, the standard *K-Means* and the standard *Spectral Clustering* algorithms are usually run several times and the best (according to some criteria) among the achieved results, is taken as the clustering solution, as the algorithms are non-deterministic in nature. Choosing a number of times to repeat these algorithms so that a good result is obtained, is generally hard. In contrast, *SilHAC* is deterministic. The hybrids have non-deterministic action due to the component non-deterministic clustering algorithm which only corresponds to the bottom level of the clustering hierarchy. Subsequent steps of building the hierarchy are deterministic. So the chance of getting bad results are comparatively less in the case of *SilHAC* and the hybrids compared to the *SC-SI* and *KM-SI* pipelines.

Furthermore, there are massive savings in execution time for the proposed hybrids when compared to the traditional ‘*clustering algorithm - internal cluster validity index*’ pipelines. For instance, consider the case of the GEO4 dataset which has 763 samples. To identify the number of clusters and the associated clustering, the *SC-SI* pipeline (with *Spectral Clustering* repeated 20 times each for *k* values ranging from 2 to 20) took close to *two hours* on a contemporary laptop computer while the *SilHAC-SC* hybrid (which initially run *Spectral Clustering* to get 25 clusters and then applied *SilHAC* on those clusters) finished the job within *seven* minutes. The bar chart in Figure 2 presents the relative execution time of *SilHAC-SC*, when the *SC-SI* pipeline is considered to take *one* unit of time to execute. Similarly, the bar chart in Figure 3 presents the relative execution time of *SilHAC-KM* when the *KM-SI* pipeline is considered to take *one* unit of time to execute. The data presented are with respect to the time taken for the results presented in Table III.

**Fig. 2.**
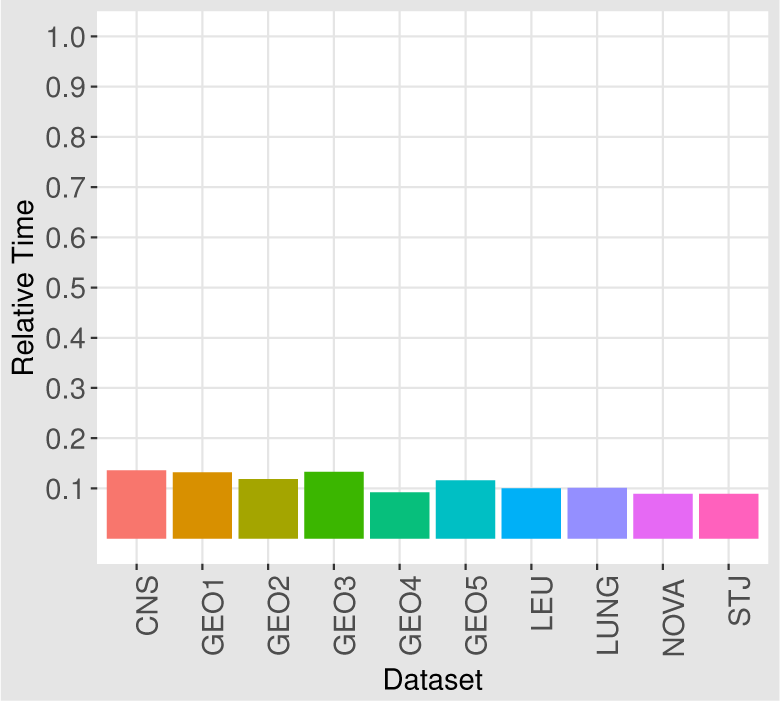
Relative execution time of *SilHAC-SC* when *SC-SI* pipeline is considered to take one unit of time to execute. The data presented are with respect to the time taken for the results shown in Table III.

**Fig. 3.**
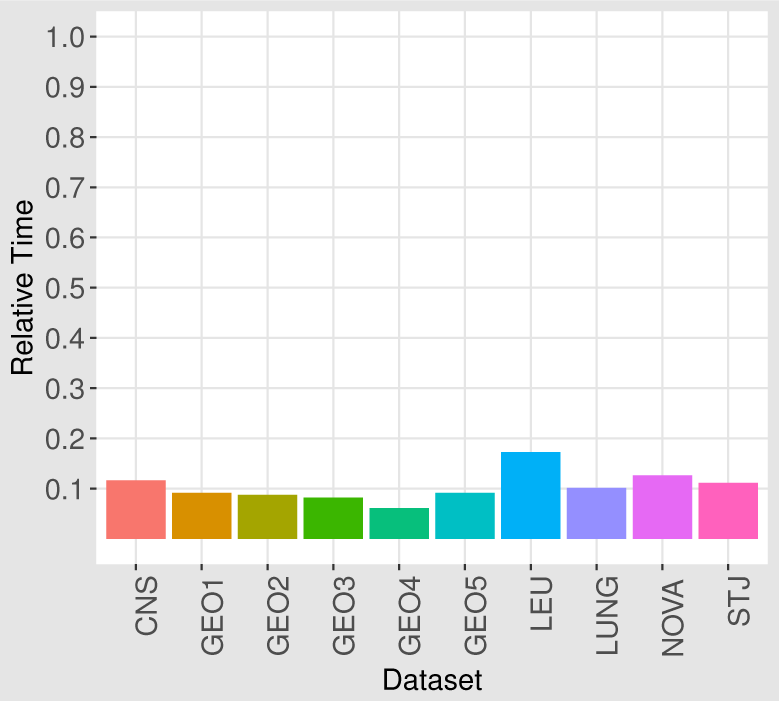
Relative execution time of *SilHAC-KM* when *KM-SI* pipeline is considered to take one unit of time to execute. The data presented are with respect to the time taken for the results shown in Table III.

A noteworthy point in the results presented in Table III is that whenever the number of clusters identified by the methods is less than the ground truth number of clusters, there is a significant increase in the *average Silhouette Index* for the clustering solution found out by the methods. The reason for this phenomenon is that *Silhouette Index* tends to prefer a lower number of clusters when there is a large variance in the inter-cluster distances. For instance, consider the dataset given in Figure 1. Clusters G2 and G3 are closer to each other and their second nearest neighbour cluster G4 is far away. Consider the calculations of *Silhouette Index* of the data points in G2 and G3 (according to Formula 1). *bi*’s of data points G2 is computed with respect to their distances to the data points in G3 and vice versa. But when G2 and G3 together are treated as one cluster, *bi*’s of the data points are computed with respect to their distances to the data points in G4 which are quite far. Definitely, there is an increase in the *ai*’s of data points of both G2 and G3 when we treat G2 and G3 are merged into one cluster but, the increase in *bi*’s is relatively much higher. Because of this, the *Silhouette Index* of data points in G2 and G3 are higher in the merged scenario than those of the data points when G2 and G3 are treated as separate clusters. The same effect may be there when we treat the groups G2, G3 and G4 together forming one cluster and G1 forming another cluster. So the methods for cluster number assessment based on *average Silhouette Index* may find *two* as the optimal number of clusters for the dataset, despite the fact that there are *four* natural groups. This is a drawback of *Silhouette Index*.

## IV. CONCLUSION

Discovery of novel subtypes of cancers leads to more refined and targeted development of drugs and which would ultimately lead to better survival rates of patients. An extensive collection of molecular data of cancer tissues are now available at publicly accessible repositories such as NCBI Gene Expression Omnibus. Cancer subtype discovery from such molecular data is an exploratory analysis, and clustering algorithms have been used extensively for the purpose. Discovery of subtypes aims at finding natural groups among tissue samples based on molecular data. A general method which is employed by many researchers to identify novel subtypes of cancers from such *omics* data is to use some clustering algorithm together with an *internal cluster validity index*.

In this paper, we presented a Hierarchical Agglomerative Clustering algorithm namely *SilHAC*. The algorithm employs Ward’s framework for Hierarchical Agglomerative Clustering and uses *average Silhouette Index* as the objective function to select the pair of clusters to be merged at each step. We also proposed a hybrid framework with *SilHAC* as one component where the second component can be any other clustering algorithm. The hybrids take a significantly lesser amount of time for execution than that of *SilHAC*.

A notable advantage of *SilHAC* and the hybrids, when compared to the other Hierarchical Agglomerative Clustering algorithms, is that they automatically find out the *natural* number of clusters. The number of clusters is straightforwardly indicated by the level of the clustering hierarchy which has the maximum *average Silhouette Index*. *SilHAC* achieves it without taking any input other than the dataset whereas a hybrid takes additional inputs demanded by the component clustering algorithm to attain the same. Furthermore, more reliable results are produced by *SilHAC* and its hybrids when compared to those of the traditional ‘*clustering algorithm - cluster validity index*’ pipelines constituted by non-deterministic clustering algorithms such as *K-Means* and *Spectral Clustering*.

The ability of the methods to automatically estimate the number of natural clusters in a dataset was tested on a set of five cancer gene expression datasets from *NCBI GEO* and a set of five benchmark cancer gene expression datasets. The methods were able to find reasonable estimates for the *natural* number of clusters in these datasets. Hence they can be better substitutes for the currently used *Silhouette Index* based methods to discover cancer subtypes from *omics* data.

*SilHAC* has a limitation that it takes a considerable amount of time for execution for large datasets. The hybrids of *SilHAC* solve this problem. A parallelized implementation of *SilHAC* to make a more scalable version of it is a future work considered.

Moreover, *Silhouette Index* is found to have a shortcoming that it tends to prefer a lower number of clusters especially in datasets where there is considerable variation in the inter-cluster distances. This variation influences the *Silhouette Index* based cluster number identification methods to prefer lower numbers of clusters than actual. Rectifying this drawback of *Silhouette Index* is another line of future research.

## REFERENCES

[1] Marcilio CP de Souto, Ivan G Costa, Daniel SA de Araujo, Teresa B Ludermir, and Alexander Schliep, “Clustering cancer gene expression data: a comparative study,” BMC bioinformatics, vol. 9, no. 1, pp. 497, 2008.

[2] Pang Ning Tan, Michael Steinbach, and Vipin Kumar, Introduction to Data Mining, (First Edition), Addison-Wesley Longman Publishing Co., Inc., Boston, MA, USA, 2005.

[3] P J Rousseeuw, “Silhouettes: A graphical aid to the interpretation and validation of cluster analysis,” J. Comput. Appl. Math., vol. 20, pp. 53–65, 1987.

[4] Todd R Golub, Donna K Slonim, Pablo Tamayo, Christine Huard, Michelle Gaasenbeek, Jill P Mesirov, Hilary Coller, Mignon L Loh, James R Downing, Mark A Caligiuri, et al., “Molecular classification of cancer: class discovery and class prediction by gene expression monitoring,” Science, vol. 286, no. 5439, pp. 531–537, 1999.

[5] Christian Wiwie, Jan Baumbach, and Richard Ro¨ttger, “Comparing the performance of biomedical clustering methods,” Nature methods, vol. 12, no. 11, pp. 1033, 2015.

[6] Menglan Cai and Limin Li, “Subtype identification from heterogeneous tcga datasets on a genomic scale by multi-view clustering with enhanced consensus,” BMC Medical Genomics, vol. 10, no. 4, pp. 75, Dec 2017.

[7] Dingming Wu, Dongfang Wang, Michael Q. Zhang, and Jin Gu, “Fast dimension reduction and integrative clustering of multi-omics data using low-rank approximation: application to cancer molecular classification,” BMC Genomics, vol. 16, no. 1, pp. 1022, Dec 2015.

[8] Qianqian Shi, Chuanchao Zhang, Minrui Peng, Xiangtian Yu, Tao Zeng, Juan Liu, and Luonan Chen, “Pattern fusion analysis by adaptive alignment of multiple heterogeneous omics data,” Bioinformatics, vol. 33, no. 17, pp. 2706–2714, 2017.

[9] Prabhakar Chalise and Brooke L. Fridley, “Integrative clustering of multi-level omic data based on non-negative matrix factorization algorithm,” PLOS ONE, vol. 12, no. 5, pp. 1–18, 05 2017.

[10] Zongcheng Li, Yaowen Chen, Shuofeng Hu, Jian Zhang, Jiaqi Wu, Wu Ren, Ningsheng Shao, and Xiaomin Ying, “Integrative analysis of protein-coding and non-coding rnas identifies clinically relevant subtypes of clear cell renal cell carcinoma,” Oncotarget, vol. 7, no. 50, pp. 82671, 2016.

[11] Bo Wang, Aziz M Mezlini, Feyyaz Demir, Marc Fiume, Zhuowen Tu, Michael Brudno, Benjamin Haibe-Kains, and Anna Goldenberg, “Similarity network fusion for aggregating data types on a genomic scale,” Nature methods, vol. 11, no. 3, pp. 333, 2014.

[12] Joe H Ward Jr, “Hierarchical grouping to optimize an objective function,” Journal of the American statistical association, vol. 58, no. 301, pp. 236–244, 1963.

[13] Paola Tellaroli, Marco Bazzi, Michele Donato, Alessandra R Brazzale, and Sorin Drăghici, “Cross-clustering: A partial clustering algorithm with automatic estimation of the number of clusters,” PloS one, vol. 11, no. 3, pp. e0152333, 2016.

[14] R Core Team, R: A Language and Environment for Statistical Computing, R Foundation for Statistical Computing, Vienna, Austria, 2017.

[15] Martin Maechler, Peter Rousseeuw, Anja Struyf, Mia Hubert, and Kurt Hornik, cluster: Cluster Analysis Basics and Extensions, 2017, R package version 2.0.6 — For new features, see the ‘Changelog’ file (in the package source).

[16] Alexandros Karatzoglou, Alex Smola, Kurt Hornik, and Achim Zeileis, “kernlab – an S4 package for kernel methods in R,” Journal of Statistical Software, vol. 11, no. 9, pp. 1–20, 2004.

[17] N Nidheesh, KA Abdul Nazeer, and PM Ameer, “An enhanced deterministic k-means clustering algorithm for cancer subtype prediction from gene expression data,” Computers in Biology and Medicine, vol. 91, pp. 213–221, 2017.

[18] Tanya Barrett, Stephen E Wilhite, Pierre Ledoux, Carlos Evangelista, Irene F Kim, Maxim Tomashevsky, Kimberly A Marshall, Katherine H Phillippy, Patti M Sherman, Michelle Holko, et al., “NCBI GEO: Archive for functional genomics data sets update,” Nucleic Acids Research, vol. 41, no. D1, pp. D991–D995, 2012.

[19] Stefano Monti, Pablo Tamayo, Jill Mesirov, and Todd Golub, “Consensus clustering: A resampling-based method for class discovery and visualization of gene expression microarray data,” Machine Learning, vol. 52, no. 1, pp. 91–118, 2003.

[20] Banumathy Gowrishankar, Christopher G Przybycin, Charles Ma, Subhadra V Nandula, Brian Rini, Steven Campbell, Eric Klein, RSK Chaganti, Cristina Magi-Galluzzi, and Jane Houldsworth, “A genomic algorithm for the molecular classification of common renal cortical neoplasms: development and validation,” The Journal of Urology, vol. 193, no. 5, pp. 1479–1485, 2015.

[21] Andrea M Griesinger, Rebecca J Josephson, Andrew M Donson, Jean M Mulcahy Levy, Vladimir Amani, Diane K Birks, Lindsey M Hoffman, Steffanie L Furtek, Phillip Reigan, Michael H Handler, et al., “Interleukin-6/STAT3 pathway signaling drives an inflammatory phenotype in Group A ependymoma,” Cancer Immunology Research, pp. canimm–0061, 2015.

[22] Florence MG Cavalli, Marc Remke, Ladislav Rampasek, John Peacock, David JH Shih, Betty Luu, Livia Garzia, Jonathon Torchia, Carolina Nor, A Sorana Morrissy, et al., “Intertumoral heterogeneity within Medulloblastoma subgroups,” Cancer Cell, vol. 31, no. 6, pp. 737–754, 2017.

[23] Anna Karlsson, Hans Brunnstro¨m, Patrick Micke, Srinivas Veerla, Johanna Mattsson, Linnea La Fleur, Johan Botling, Mats Jo¨nsson, Christel Reuterswa¨rd, Maria Planck, et al., “Gene expression profiling of large cell lung cancer links transcriptional phenotypes to the new histological WHO 2015 classification,” Journal of Thoracic Oncology, 2017.

